# Functional autophagy gene set signature and state classification reveal a link between autophagy induction, lysosomal activity, and poor prognosis in glioblastoma

**DOI:** 10.1101/2025.09.08.674856

**Authors:** Camilla Brückmann de Mattos, Daphne Torgo, Eduardo C. Filippi-Chiela, Guido Lenz

## Abstract

Autophagy is an essential mechanism for maintaining cell homeostasis and, when dysregulated, is related to various pathologies. In cancer, it functions as a double-edged sword, either as a tumor suppressor or as a tumor promoter. Elevated GFP-LC3 puncta and increased levels of autophagy-related proteins, commonly interpreted as markers of high autophagic activity, are often associated with poor prognosis in tumors. However, current assessments of autophagy largely rely on isolated cellular and protein-level observations. While transcriptomic approaches offer broader insights, they often utilize large gene signatures without adequately accounting for the dynamic nature of autophagic flux. In this study, we propose a functional gene set signature for autophagy based on the rational mapping of flux stages, enabling the inference of distinct autophagy states. Using a supervised machine learning approach, we developed two indices - AutoIndex and LysoIndex - that effectively distinguish between cells treated with autophagy inducers and inhibitors. Applying this framework, we analyzed autophagy states across four cancer types, comparing tumor samples to matched normal tissues and evaluating their association with patient survival. Among these, glioblastoma (GBM) exhibited the strongest correlation between inferred autophagy states and clinical outcomes. GBM samples characterized by high LysoIndex and low AutoIndex—corresponding to a transcriptomic induction state—were linked to significantly poorer survival. Compared to lower-grade gliomas, GBM showed elevated lysosomal activity, which may contribute to its more aggressive behavior. Additionally, GBM cell lines treated with temozolomide displayed signatures consistent with autophagy induction, supporting our model’s predictive relevance. These results reinforce the understanding and monitoring of autophagy, and suggest that, in the case of GBM, its negative influence may represent a potential target for novel therapeutic interventions.

## Introduction

As extensively described in the literature, macroautophagy – hereafter referred to as autophagy – is a fundamental catabolic pathway conserved from yeast to mammals. It is the basis for recycling damaged cellular components and is essential for maintaining cellular homeostasis. The main function of autophagy is to sense metabolic stress, engulfing cytoplasmic components and directing them to the lysosome, where they are degraded by hydrolytic enzymes [1–3]. The autophagic flux consists of five main steps: initiation (a), nucleation (b), elongation (c), maturation (d) and degradation (e). The initiation (a) involves the sensing of metabolic stress, triggering the *de novo* assembly of autophagosomes. Next, the phagophore assembly site (PAS) is established during the nucleation (b) and evolves from a phagophore (PG) to a closed autophagosome through the elongation step (c). During maturation (d), autophagosomes fuse with lysosomes to form autolysosomes. Finally, the sequestered material is enzymatically degraded in the degradation step (e) [1,3,4]. This flux is highly regulated by a signaling pathway, which involves the autophagy related genes (ATGs) proteins and their interactors. Initiation (a) is controlled by nutrient-sensing complexes— particularly mTORC1, which, upon sensing nutrient availability, phosphorylates and inhibits the ULK1/2 complex. Under stress, mTORC1 is inactivated, allowing ULK1/2 activation and autophagy initiation. Then three main components mediate autophagosome nucleation (b) and elongation (c): PI3K complex (b), ATG12-ATG5-ATG16 complex (c) and lipidated LC3 (LC3-II) associated with SQSTM1 (c). Following, the maturation (d) step involves lysosomal proteins – VAMP8, STX7, LAMP and RAB7 – as well as autophagosomal proteins – LC3, STX17 and YKT6 [1,3–5].

Because each step is so tightly regulated, any dysfunction can have profound cellular consequences, impairing important regulatory roles played by autophagy, such as cell integrity protection mechanisms from stress controlling to cell fate decisions [6], being the cause of different pathologies. One of them – and the first to be related to autophagy malfunctions – is cancer [7]. Nevertheless, autophagy is described as a double agent in cancer malignancy, playing both tumor suppressor and promoter roles. As a tumor suppressor, autophagy can prevent the initiation of tumorigenesis by maintaining genomic stability. In contrast, as a tumor promoter, autophagy protects cells against stress induced by chemotherapy [8,9]. In the literature, it is proposed that tumors with high autophagic flux present higher malignancy, which is already described in melanoma [10] and breast cancer [11]. In addition, some studies have already shown that a negative pharmacological modulation of autophagy can enhance chemotherapy efficiency [12,13]. Therefore, the monitoring of autophagy levels and the possible modulation of this phenotype can be a prospective target to improve cancer treatment. The state of the art of monitoring autophagy involves punctual observations of the autophagy flux. Autophagic flux is related to the dynamics of autophagy, referring specifically to the rate at which cellular components are degraded within autolysosomes [14]. Some of the approaches include GFP-LC3 puncta quantification– since they can indicate the number of autophagosomes in a cell – and western blot monitoring of LC3-I, LC3-II and SQSTM1 protein levels, with and without the addition of a lysosome blocker such as hydroxychloroquine or bafilomycin 1. For a transcriptional analysis, the autophagy profile can be predicted with the mRNA levels of genes like *ATG1, ATG6, ATG7, LC3, GABARAPL1, ATG9, ATG12, ATG13, ATG14, ATG29, WIPI1* and *SQSTM* [15]. However, these approaches do not provide an integrative, multifactorial analysis capable of differing autophagy flux induction and inhibition conditions – hereafter states (i.e., transient functional conditions of the phenotype). Whole transcriptome analysis has emerged as a key tool to characterize cellular phenotypes as distinct states [16], making it a promising approach for comprehensive profiling of autophagic flux and its prognostic relevance. Many studies use large autophagy-related gene signatures from databases such as the Human Autophagy Database (HADb) and the Human Autophagy Modulator Database (HAMdb) [17–19]. However this approach may not be the most accurate to predict autophagy flux, since it includes many genes with typically altered profiles in cancer, such as *P53* and *RB1* [20]. Besides that, evaluating gene sets corresponding to distinct phases of the autophagic flux could provide a more comprehensive understanding, as compensatory mechanisms between these phases might obscure dysfunctions when assessed in isolation.

In order to enhance the characterization of autophagy flux in relevant states and their relationship with prognosis, we propose an optimized autophagy gene set signature with a functional focus, considering both autophagic flux and lysosome essential components, as well as regulatory proteins, and dividing it into relevant gene sets, which were analyzed by Gene Set Variation Analysis (GSVA). Using datasets of cells treated with autophagy modulators, we developed a separation model to predict autophagy states from expression alterations among conditions. Using this model, we identified autophagy state profiles in patient samples for four cancer types among The Cancer Genome Atlas (TCGA) projects: Glioblastoma Multiforme (GBM), Breast Invasive Carcinoma (BRCA), Lung Adenocarcinoma (LUAD) and Colon Adenocarcinoma (COAD). The cancer types with more significant correlation of differential states and prognosis were GBM and COAD, indicating that an induction state is more related to a poor prognosis. Investigating more deeply the states in GBM, we compared high-grade and low-grade (LGG project) glioma autophagy states and states induced by typical therapy. Results showed a higher incidence of induction states in high-grade glioma and an increase occupation of this state caused by chemotherapy. However, the lack of statistical significance may be explained by the limited number of samples, which could have reduced the power of the analysis. Collectively, our results propose a method to monitor autophagy via state identification and suggest that the induction state—particularly in GBM—is associated with poor prognosis.

## Results

### An autophagy gene signature can be established and divided into gene sets based on function in autophagic flux

This work comprises three main steps: gene signature establishment, autophagy states definition and states application in clinical data (Figure 1). For the gene signature, we summarized an extensive list of genes related to autophagy, focusing on its functionality within the autophagic flux. Therefore, we analyzed the traditional HADb searching specifically for ATG proteins. In addition, to define the autophagy-related genes (ARGs) for each step (initiation, nucleation, elongation and maturation), we analyzed the autophagy pathway as disposed in Kyoto Encyclopedia of Genes and Genomes (KEGG) and Reactome. With that, we identified 26 genes related to autophagy and they were divided into the four following gene sets (Figure 2): initiation (5 genes), nucleation (6 genes), elongation (8 genes) and maturation (7 genes). Besides intrinsic autophagy related genes, we included two regulatory gene sets, with the main autophagy regulators proposed in the literature [3]: Positive Regulation (4 genes, *CAMKK2, AMPK, TSC1* and *TSC2*) and Negative Regulation (2 genes, *MTOR* and *PKA*).

**Figure 1.**
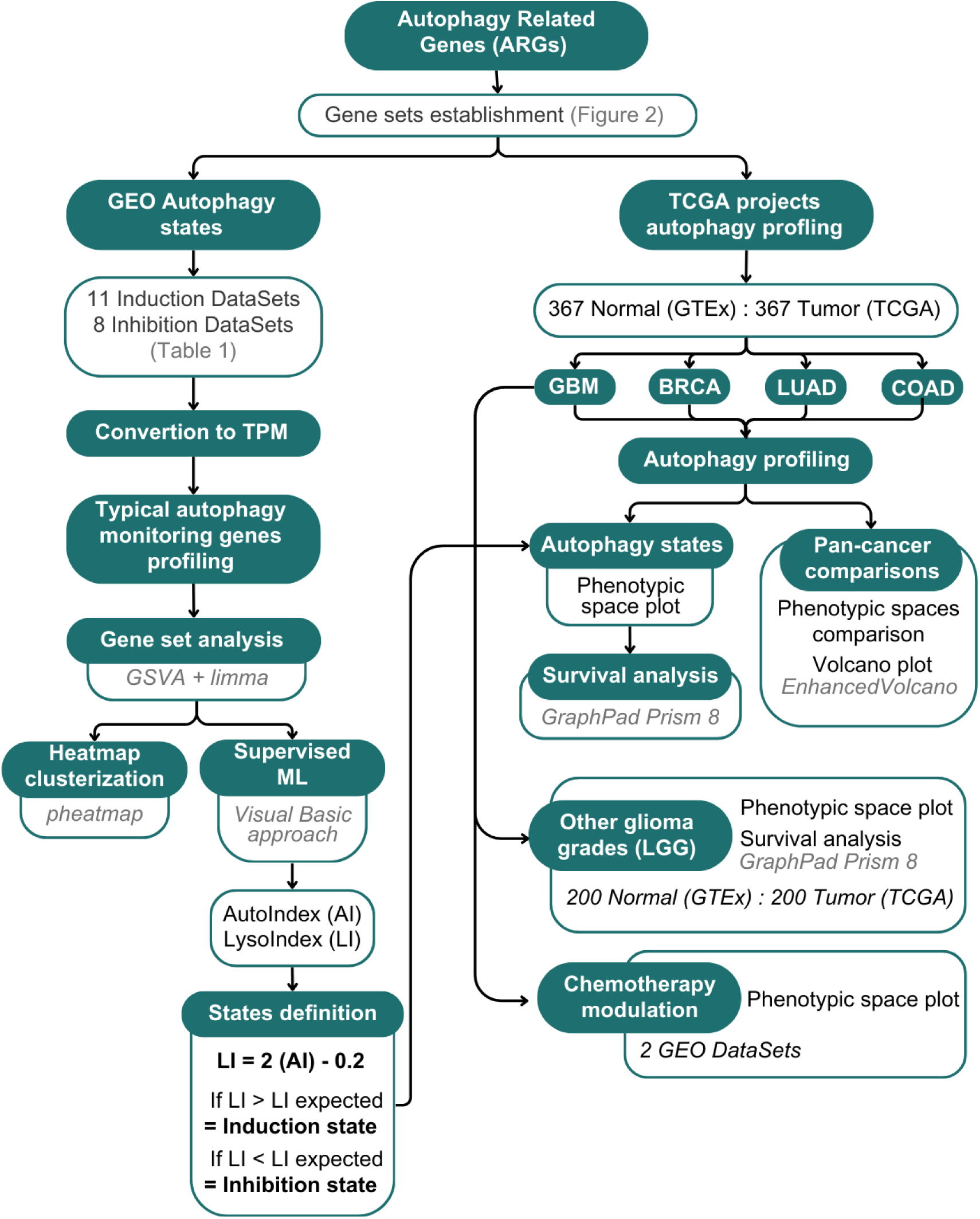
Overview of the study design and methodology.

**Figure 2.**
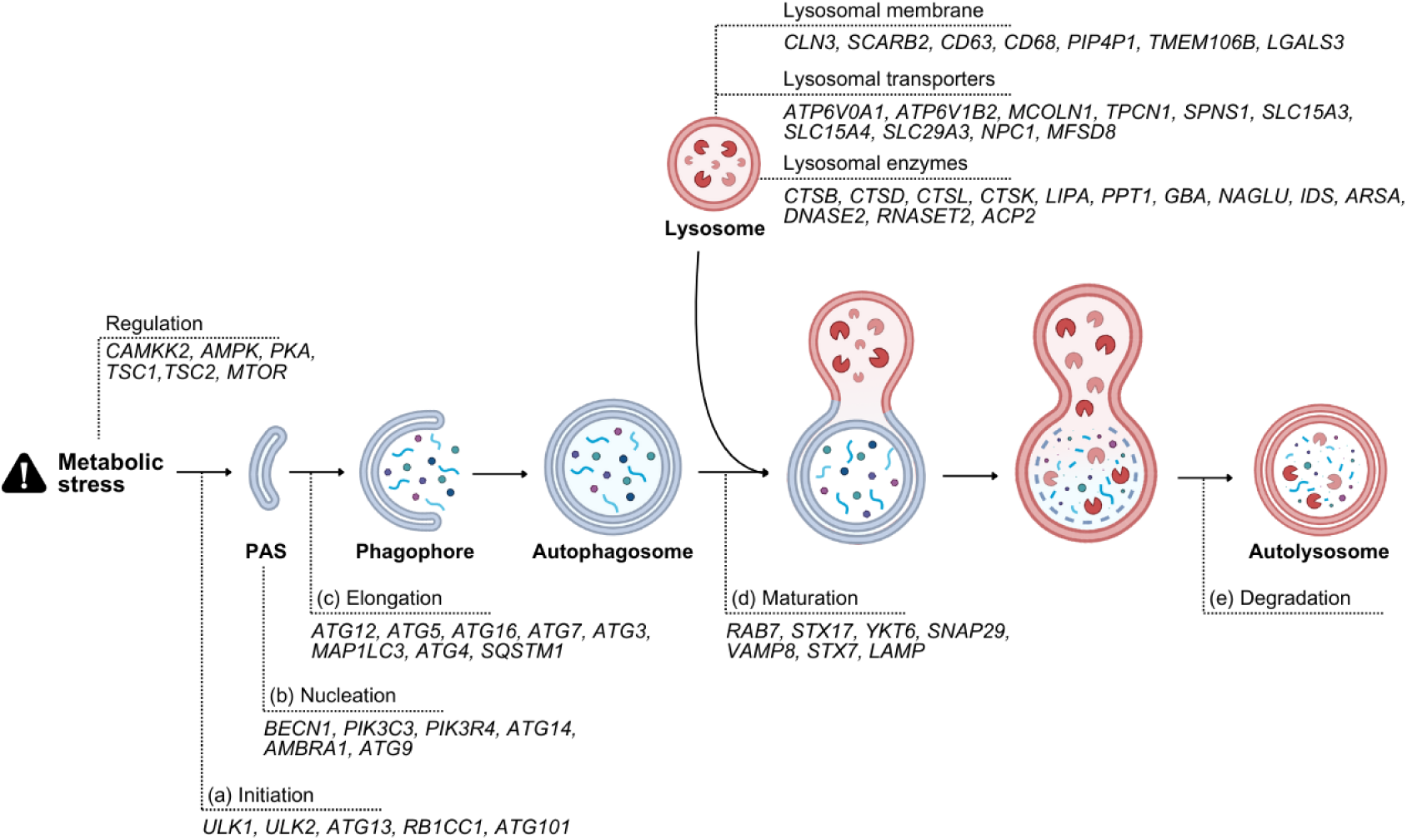
Autophagy gene signature gene sets divided among autophagic flux steps.

With this signature, we also included gene sets exclusively for lysosomal function. They were divided into three gene sets: Lysosomal membrane proteins (L. membrane), Lysosomal transporters (L. transporters) and Lysosomal enzymes (L. enzymes). As L. membrane, there were selected 7 genes (Figure 2), which are related to formation and structure of lysosomal membrane; as L. transporters, there were selected 10 genes (Figure 2), which include protons, ions, lipid and amino-acids transporters responsible for lysosomal acidification and function present in lysosomal membrane; and as L. enzymes, there were selected 13 genes (Figure 2) related to enzymes present at the lysosome. Two broader gene sets were also defined: Autophagy (Initiation, Nucleation, Elongation and Maturation) and Lysosomal activity/L. activity (L. membrane, L. transporters and L. enzymes). This resulted in a total of 62 genes that are described in Figure 2 and Table S1.

### Differential autophagic cell states can be identified among different treatment conditions

As well as being controlled by metabolic stress levels, autophagy can be pharmacologically modulated. Its induction can be mediated by cell stressors like tunicamycin and thapsigargin (endoplasmic reticulum stressors) and protein regulators like rapamycin or its analogue Everolimus (inhibitors of mTORC1). On the other hand, inhibition can be promoted by drugs that inhibit nucleation, like 3-MA, or maturation, like bafilomycin and (hydroxy)chloroquine [21,22]. Therefore, it is expected to identify cells with different autophagy states when treated with different autophagy modulators. In this sense, we selected 19 Gene Expression Omnibus (GEO) data sets with expression profiling by high throughput sequencing, hereafter bulk-RNA-seq, of cell lines exposed to autophagy modulators (Table 1, Table S2). Eleven of them included induction conditions (with exposure to rapamycin, everolimus, starvation conditions, imperatorin or Z36) and 8 of them included inhibition conditions (with exposure to chloroquine, hydroxychloroquine, bafilomycin, BRDA siRNA, spautin-1, elaiophylin or ambroxol). The main target for autophagy induction was the initiation/nucleation; and for inhibition, the initiation or maturation (Figure 3A).

**Figure 3.**
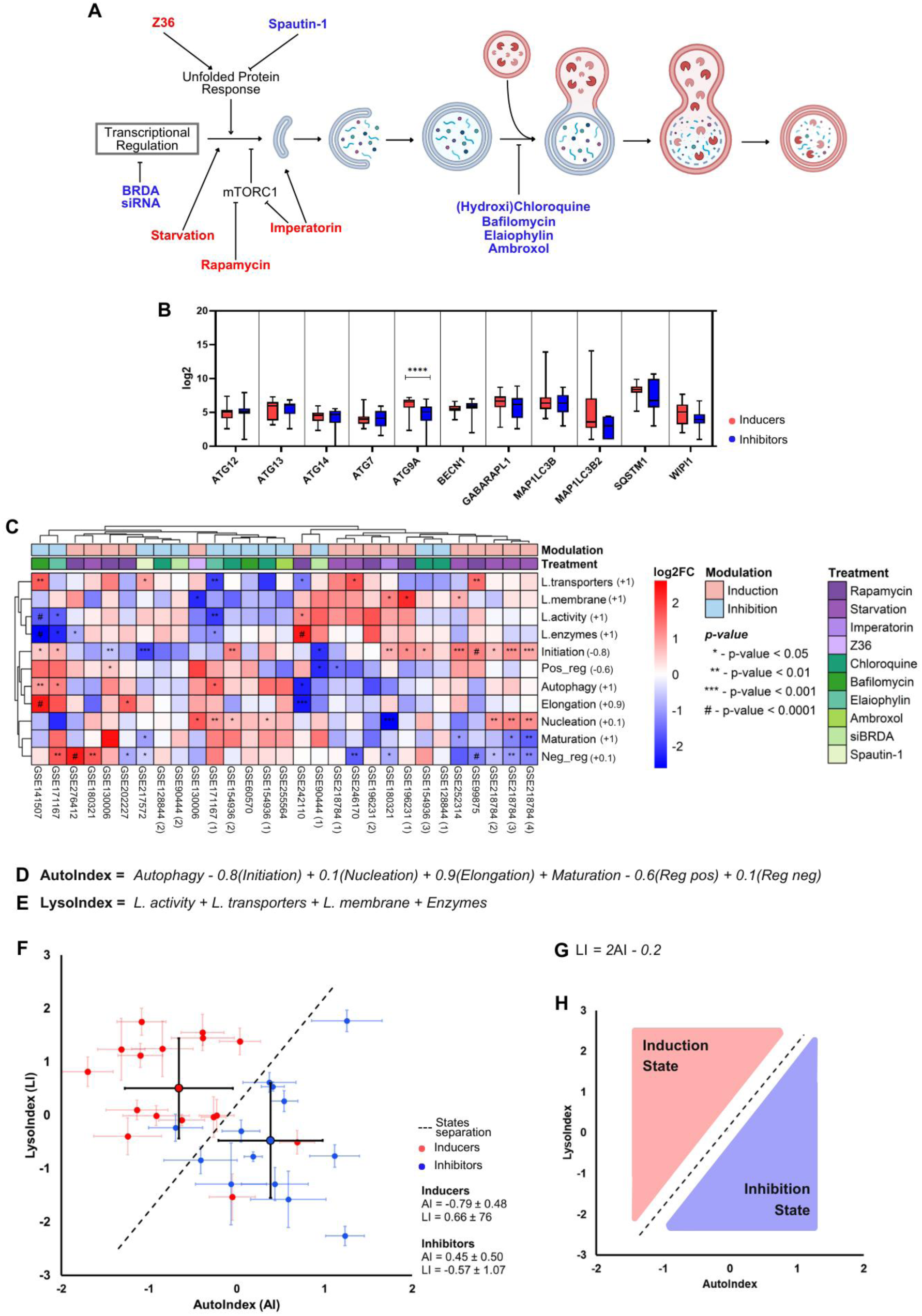
Autophagy states characterization and separation. (**A**) Autophagic flux with indication of autophagy modulators target of action. In red, inducers. In blue, inhibitors. (**B**) Expression of typical ARGs for autophagy monitoring in samples exposed to autophagy inhibitors and inducers. Multiple t-tests were carried out to compare the expression data between conditions. (**C**) Heatmap clustering of GEO samples with columns and rows clustering. Red to blue scale indicated high to low log2FC values. Differential expression analysis significance indicated by p-value annotation in each heatmap cell. Predicted modulation and treatment of each data set indicated as column annotations. Coefficient calculated by supervised ML indicated next to respective gene set name. (**D**) Proposed formula for AutoIndex calculation. (**E**) Proposed formula for LysoIndex calculation. (**F**) States plot for GEO samples. Average and dispersion values of each index indicated for each condition. (**G**) Final states separation formula. (**H**) Summary of the separation proposed by the states plot.

**Table 1.**
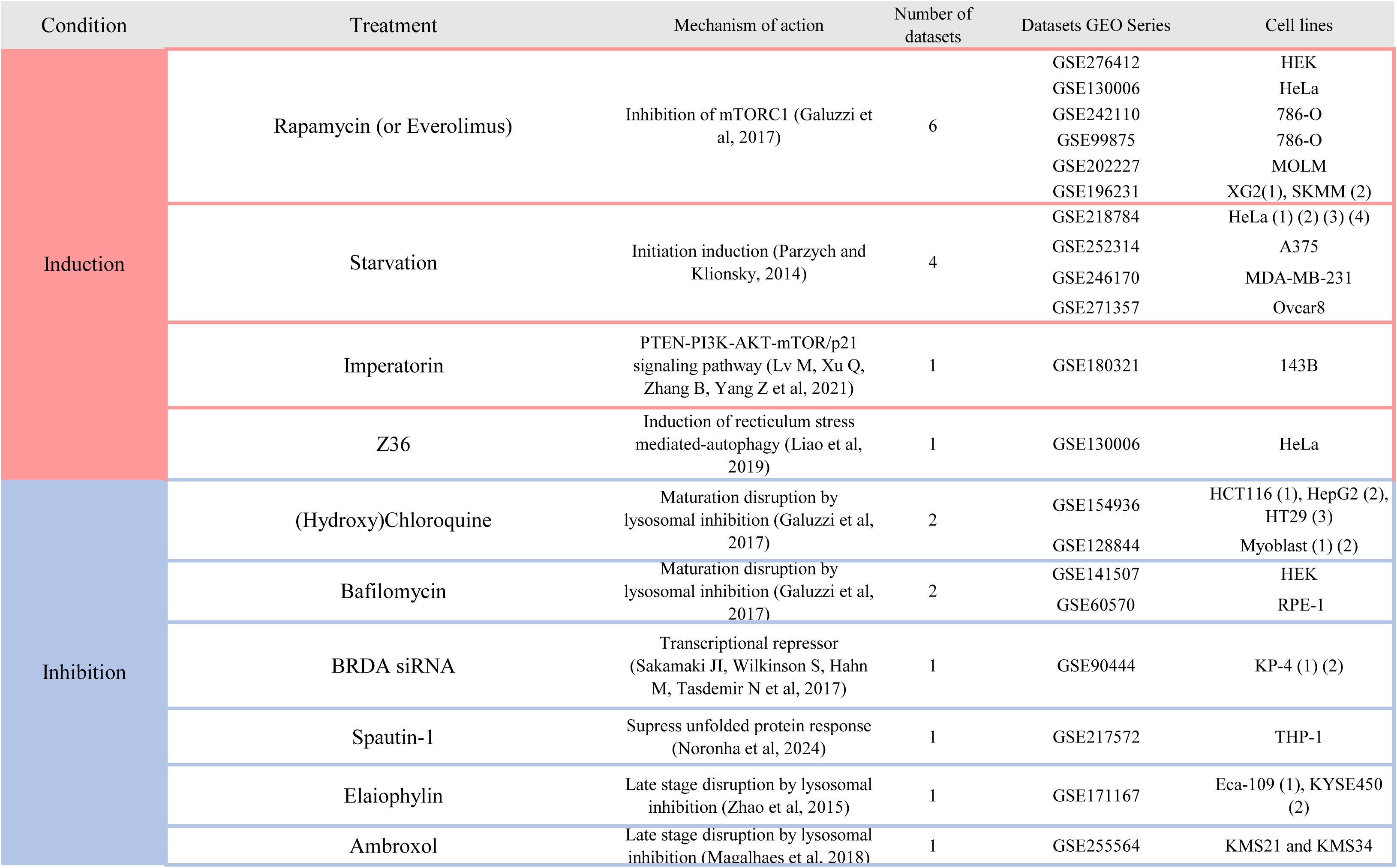
GEO data sets for autophagy states definition.

As a primary differential expression screening, normalized expression as log2 values of typical autophagy monitoring genes, as proposed by Klionsky, Abdel-Aziz, Amal Kamal, Abdelfatah and Sara et al 2021, was carried out. The comparison between distributions among samples exposed to induction and inhibition conditions showed a significant increase only in ATG9A for induction conditions (Figure 3B). To better comprehend states differences and not rely only on one differentially expressed gene, we continue the analysis using the gene signature with gene sets previously established.

The expression data was analyzed using GSVA methodology, which returned a GSVA score for each sample. Among each GEO project, these values were used to calculate differential expression of gene sets between control and treated samples. To integrate these data into meaningful autophagy flux differentiation indices, we implemented two approaches on the data sets with significant results: heatmap clustering and supervised machine learning. With the heatmap, columns and rows clustering returned a division of the samples into two main groups (Figure 3C). One was composed of data sets exposed to autophagy inducers, which showed a prevalence of positive log2FC values for lysosomal activity gene sets and negative for autophagy related ones. And the other was composed of data sets with autophagic flux inhibitors treatments, with an opposite behavior on log2FC values. Despite the overall separation, some data points failed to cluster as expected, highlighting residual overlap between groups.

Thus, we implemented a basic supervised machine learning (ML) approach that assigned weighting coefficients to be applied to the log2FC values of each gene set, aiming to optimize the separation between differential states. As a result, two index values were defined, representing the output of these weighted calculations. AutoIndex (AI) was defined as the sum of the weighted values for the gene sets related to autophagy (Initiation, Nucleation, Elongation, Maturation, Positive Regulation, and Negative Regulation). Similarly, LysoIndex (LI) was calculated as the sum of the weighted values for the gene sets associated with lysosomal function (Lysosomal Activity, Lysosomal Membrane, Lysosomal Enzymes, and Lysosomal transporters). We set the ML to prioritize low AI and high LI values for induction samples and the opposite behavior for inhibition samples, to better match the heatmap observations. The best autophagy states dividing coefficient values can be seen in Figure 3D for AI calculation and 3E for LI. The final division plot (Figure 3F) showed only 3 data sets that were not correctly divided: GSE154926 (3) (Hydroxychloroquine), GSE130006 (Z36) and GSE202227 (Rapamycin) (Figure S1A-B). The final proposed linear equation that better differs the differential autophagy conditions was *LI = 2AI - 0.2* (Figure 3G-H), where *y* stays for an expected LI value for a specific AI value, represented by *x*. When the observed LI exceeds the expected LI, the sample is interpreted as being in an induction state. Conversely, when the observed LI is lower than the expected LI, the sample is classified as being in an inhibition state. This method for distinguishing autophagy states, along with the “Phenotypic Space Plot” data visualization, was used in the subsequent analyses. An automated table supporting this approach is provided in Table S3.

### Autophagic cell states can be identified in patient samples and related to overall survival, mainly in glioblastoma

To implement the analysis of autophagy states in clinical data, we characterized the autophagy-focused phenotypic space in samples of four projects from TCGA: GBM (Glioblastoma Multiforme), BRCA (Breast Invasive Carcinoma), LUAD (Lung Adenocarcinoma) and COAD (Colon Adenocarcinoma). From each project, 367 solid tumor samples were randomly selected and a matching GTEx cohort of 367 related normal samples per cancer type was used as control (Table S4).

Phenotypic space plot indicated a significant difference in AI and LI values between tumor and control samples for all cancer types analyzed and a prevalence of autophagy flux inhibition state. Percent values of each state for tumor samples are indicated at phenotypic space plot in Figure 4A. For all cancers analyzed, AI was higher in tumor samples in relation to control whereas LI was slightly lower for all cancers, except GBM (Figure 4A). Comparing index final values among projects, GBM was the one with more increased AI and LI, differing significantly from every other cancer type (Figure 4B-C). Besides the index comparisons, we analyzed the volcano plots of the gene sets among its respective genes. Log2FC values indicates the prevalence of autophagic flux related gene sets with positive values and lysosomal activity related gene sets with negative ones, being the most notable exception the L. membrane value for GBM (Figure S2A), what correlated with the predominance of an inhibition state in all cancer types.

**Figure 4.**
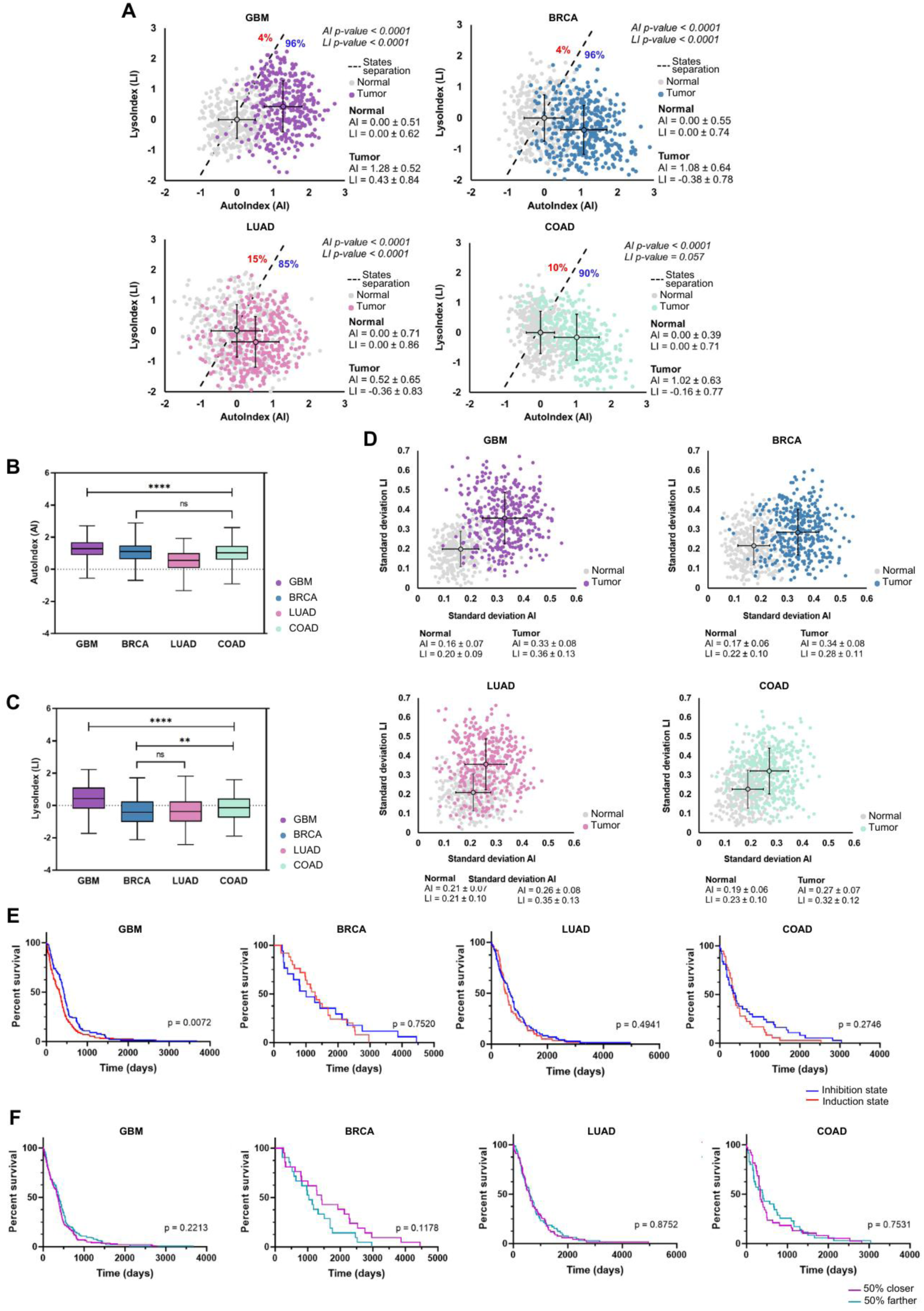
Autophagy states characterization in different cancer types. (**A**) Phenotypic space plot for TCGA Projects. Percentage of each state indicated at the graphic in red (induction) and blue (inhibition). Average and dispersion values of each index indicated for both normal and tumor samples. Mann-Whitney U test was carried out to compare distribution values of each index between conditions. Box plot of the distribution of AI (**B**) and LI (**C**) values for each cancer type. Kruskal-Wallis test was carried out to compare the multiple index values. (**D**) Standard deviation plot of values that comprise each index. Mean standard deviation values and its deviation indicated for both normal and tumor samples. Kaplan-Meier survival analysis for each cancer type comparing autophagy states (**E**) and distance from normal samples (**F**). P-values are indicated.

Given that each index was derived from multiple gene set values, we assessed the internal variability within each index to understand their heterogeneity. Evaluating heterogeneity in cancer is crucial for comprehending the complexity of various phenotypes and their influence on treatment responses. [23]. To this end, we calculated the standard deviation of each index – based on its associated gene sets - for every sample. As expected, the standard deviation plots revealed greater heterogeneity in tumor samples compared to normal tissues (Figure 4D). Specifically, the AutoIndex exhibited higher variability in BRCA and GBM, while the LysoIndex showed increased heterogeneity in GBM and LUAD. These findings underscore the pronounced heterogeneity of GBM, a characteristic often associated with its malignancy [24].

To expand the analysis of autophagy states to clinical implications, we conducted a series of Kaplan-Meier analysis. Given the limited number of samples classified within the induction state, we adjusted the separation threshold to intersect the median point, thereby dividing the dataset into two equal halves (50% above and 50% below). To preserve the original state characteristics, this new threshold line was drawn parallel to the initial state-separation boundary (Figure S2B). The resulting Kaplan-Meier curves indicated a significant survival difference between induction and inhibition state only for GBM patient samples, being the induction state the one with the poorest prognosis (Figure 4F). When selected only the patient that had an overall survival of at least 300 days, the COAD samples also showed a significant difference with the same behavior observed at GBM (Figure S2D). To assess whether the autophagy phenotypic space can predict significant differences in survival solely based on the induction–inhibition axis, we also tested whether the normal–tumor axis had any impact on survival. However, no Kaplan-Meier curve showed significant differences between groups (Figure 4G). One last alternative division was implemented, separating samples into terciles, likewise previously done with the median (Figure S2C). Again, only GBM samples showed a significant difference among groups corroborating to the conclusion that the higher the AI value and the lower the LI value, the better the prognosis for glioblastoma, which cannot be seen in other cancer types. Since GBM was the cancer type with more conclusive results, it will be the focus of the following analysis.

### The correlation of autophagic states and prognosis is more significant to high glioma grades, which show increased lysosomal activity

To better elucidate the autophagy states in GBM, we analyzed if the differences in AI values and LI values were related to the intrinsic differential aspects of neuronal malignant tissue or were a consequence of GBM aggressive phenotype. As a comparison, samples from the other two glioma grades (G2 and G3) were obtained from TCGA project Low Glioma Grade (LGG) (Table S4). Randomly, 200 patient samples of each grade were chosen and as control, 200 samples from the previous normal brain tissue cohort were randomly selected.

Phenotypic space plot indicated an almost 100% inhibition state prevalence in both low-grade gliomas (Figure 5A). In G2 patient samples, the mean AI value was 1.36 and the mean LI value was −0.47, both significantly different from normal samples. In G3 patient samples, the mean AI value was 1.58 and the mean LI value was −0.21, where only AI value was significantly different from normal samples. When comparing G2, G3 and G4 (previous GBM analyzed samples), all indices differ significantly between glioma grades with a notable difference of the LI mean values of G4 to the other subtypes (Figure 5B). Interestingly, the standard deviation plots also reveal that G2 and G3 are much more homogeneous in terms of LI values, showing levels of variability similar to those of normal samples (Figure 5C). In contrast, for AI, G2 and G3 appear more heterogeneous when compared to both G4 and normal tissues (Figure 5D). Analyzing these differences beyond index values, gene set differential expression showed Autophagy and Maturation significant log2FC positive values in both G2 and G3 (Figure S3A) whereas G4 included Maturation, L. membrane and Elongation (Figure S2A). The distribution of gene set log2FC values for each patient sample within its glioma subtype indicated that expression profile of this autophagy gene set signature tended to be more similar between G4 and G3 or G3 and G2 than between G2 and G4, what can be an indicative of the relationship of autophagy behavior and tumor malignancy (Figure 5E-G). The gene sets that highlight G4 differential autophagy profile – and do not differ among the other two grades - are Elongation (up-regulated), Maturation (down-regulated) (Figure 5E), L. transporters (down-regulated) (Figure 5F) and Negative regulation (down-regulated) (Figure 5G).

**Figure 5.**
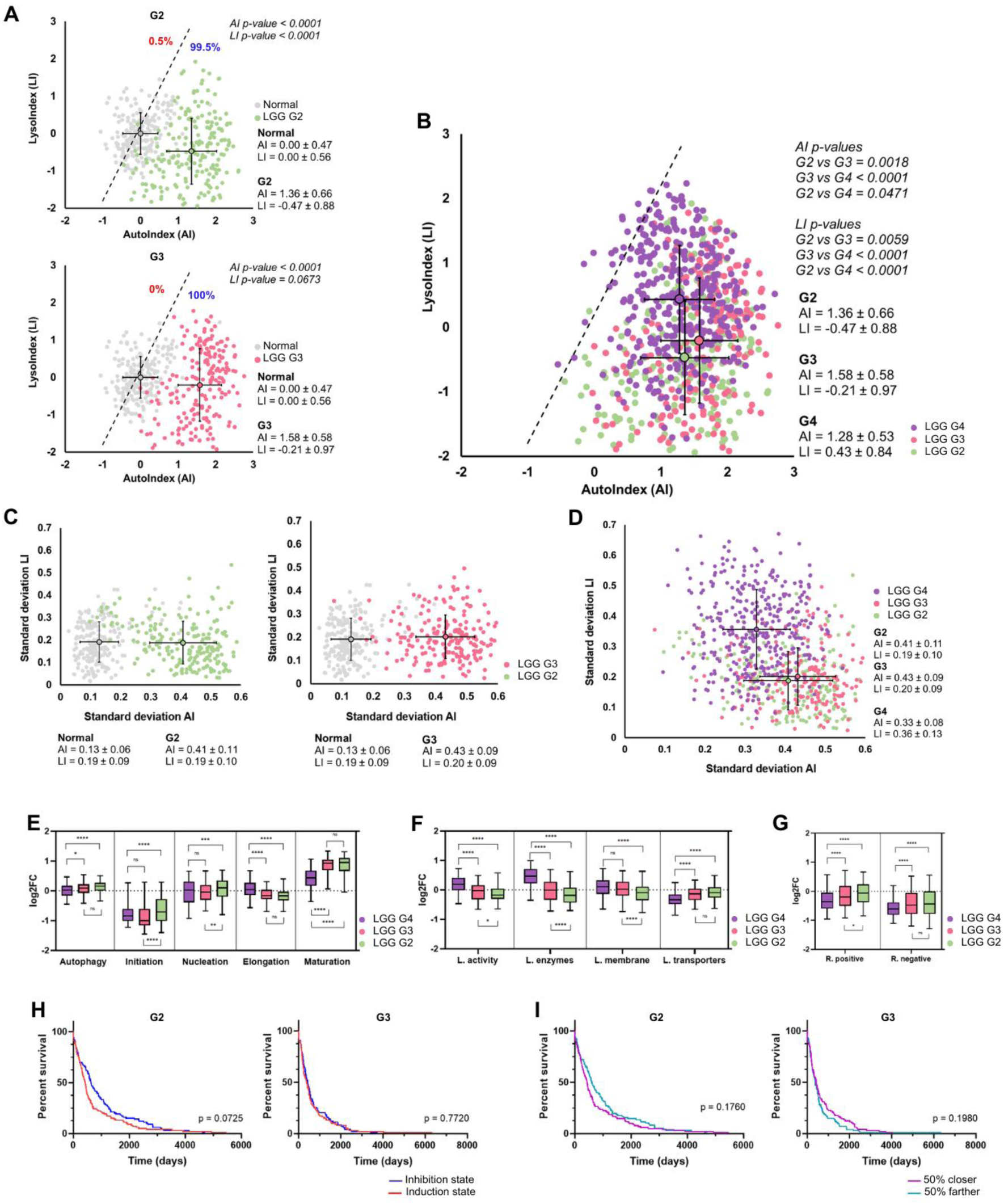
Autophagy states differentiation between different glioma grades. (**A**) Phenotypic space plot for G2 and G3 gliomas from LGG Project. Average and dispersion values of each index indicated for both normal and tumor samples. Percentage of each state indicated at the graphic in red (induction) and blue (inhibition). Mann-Whitney U test was carried out to compare distribution values of each index between conditions. (**B**) Phenotypic space plot comparing G2, G3 and G4 glioma samples states distributions. Average and dispersion values of each index indicated for all conditions. Kruskal-Wallis test was carried out to compare the multiple index values. Standard deviation plot of values that comprise each index for individual grades (**C**) and all grades together (**D**). Mean standard deviation values and its deviation indicated for both normal and tumor samples. Box plot of the distribution of autophagic flux related gene sets (**E**), lysosomal activity related gene sets (**F**) and regulation gene sets (**G**) log2FC values for each glioma grade. ANOVA test was carried out to compare expression between grades. Kaplan-Meier survival analysis for each low glioma grade comparing autophagy states (**H**) and distance from normal samples (**I**). P-values are indicated.

In sense of clinical implications, it was expected higher overall survival rates at low grades glioma [25]. However, we aimed to explore whether the autophagy states would also influence LGG survival as shown in GBM samples. Therefore, the median separation (Figure S3B) and distance to normal samples were also employed here. The resulting Kaplan-Meier curves indicated no significant differences between groups for the two analyses in both G2 and G3 glioma samples (Figure 5F-G). These results suggest that the increase in the LI index as well as specific gene set alterations in G4 samples might be an indicative of a differential autophagy profile of GBM tumors which, as seen, can be related to poor prognosis.

### TMZ treatment can alter the autophagic states in glioma samples and cell lines

As previously reported in the literature, temozolomide (TMZ) treatment can enhance autophagic flux *in vitro*, which was observed by western blot and immunocytochemistry assays [26]. To investigate whether this could also be observed at the transcriptional level, we characterized the autophagy states with the proposed method of the GEO data sets GSE249544 and GSE229600. The first one contains bulk RNA-seq data from patient-derived glioblastoma cell lines exposed to radiation and TMZ. Phenotypic space plot showed a differential distribution of TMZ treated and radiation treated samples. For TMZ samples, the mean AI value was 0.22 and the mean LI value was 0.38, although both did not differ from untreated samples (Figure 6A). TMZ treated samples indicated 42% of samples at the induction state, the highest induction percentage among all the analyses. For radiation samples, the mean AI value was 0.66 and the mean LI value was −0.24, only AI differed significantly from control samples (Figure 6A). The percentage of autophagy induction state was 17% among radiation treated samples (Figure 6B). When compared, both treatment conditions did not significantly differ (Figure 6C). Although the index differences might be interesting, considering the presence of more induction state after treatment, they were not statistically significant, probably due to small samples sizes.

**Figure 6.**
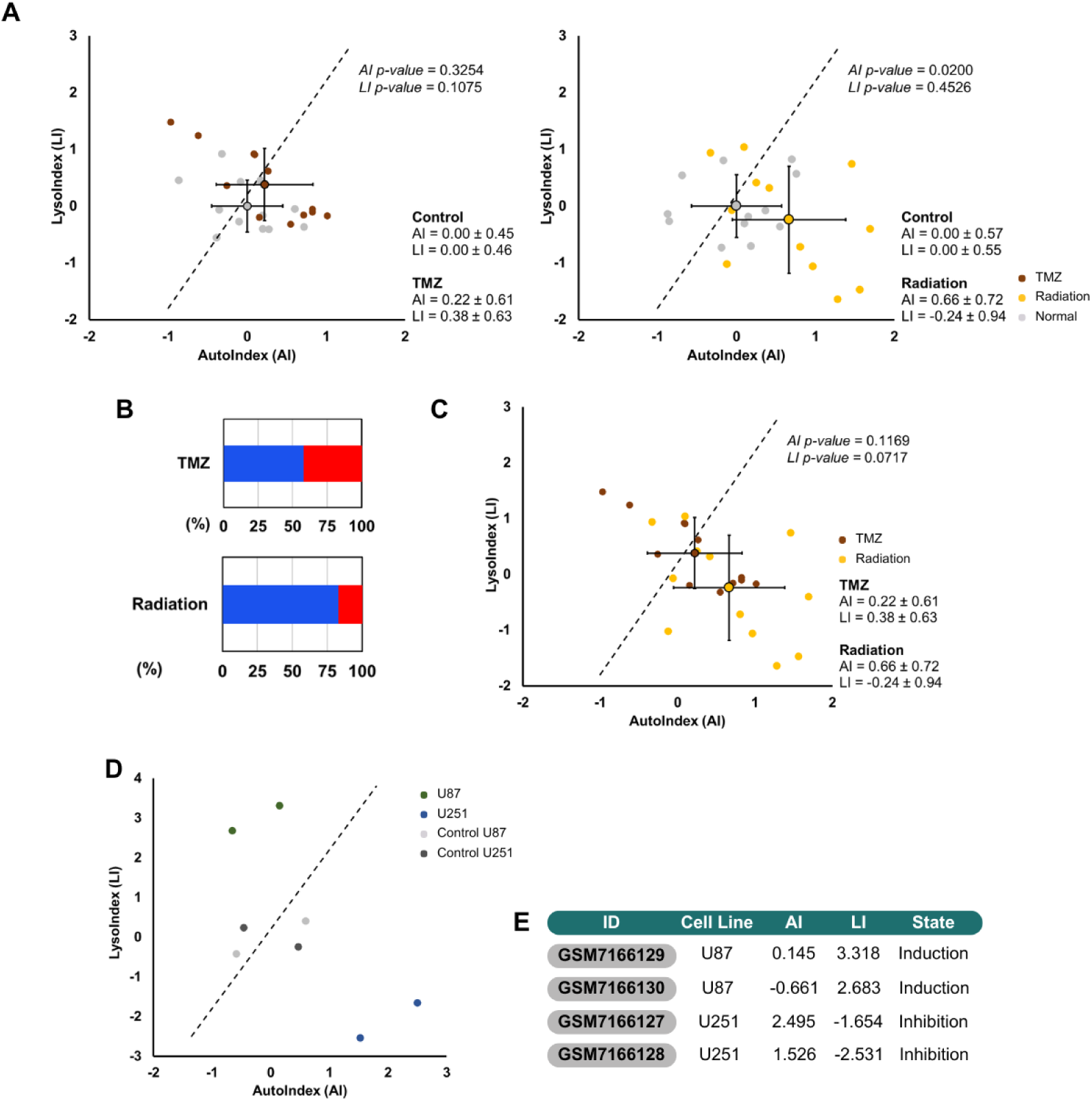
Autophagy states characterization after treatment. (**A**) States plot for TMZ and Radiation treated samples from GEO data set GSE249544. Average and dispersion values of each index indicated for both normal and tumor samples. Unpaired t-test was carried out to compare distribution values of each index between treated and control conditions. (**B**) Bar plots of states percentage for each treatment. (**C**) State plot comparing treatment states distributions. Average and dispersion values of each index indicated for both conditions. Unpaired t-test was carried out to compare distribution values of each index between treatment conditions. (**D**) States plot for TMZ treated cell samples (U87 and U251) from GEO data set GSE229600. (**E**) State data describing each bulk RNA-seq cell sample.

The second GEO data set comprises bulk RNA-seq data of two glioma cell lines treated with TMZ and respective controls. The human glioblastoma cell lines U87 and U251 exhibited different autophagy states following treatment: U87 predominantly occupied an induction state, whereas U251 showed an inhibition state. These results suggest a heterogeneous response to treatment, indicating that it may be cell type dependent. The individual index values of each sample and consequent autophagy state are described in Figure 6E. Although autophagy states analysis of treated samples did not have a great statical significance, they possibly indicate autophagy states alterations due treatment that could be further and better characterized including more data sets to this analysis.

## Discussion

Autophagy is an essential cellular process that regulates cell homeostasis. When impaired, it can be related to several pathologies, such as cancer. In tumors, autophagy can play two main but opposite roles: tumor suppressor, by maintaining genomic stability, and tumor promoter, by being a cytoprotective mechanism against therapy. In this sense, the literature already corelates tumors with poor prognosis with an augmented autophagic flux. However, the actual autophagy monitoring typical methods involve punctual observation of specific autophagy features without considering it systematically integrated with other subtle changes in the phenotype. As a complementary approach, transcriptomic analyses of autophagy represent a more accessible strategy, given the availability of numerous open-access databases, and offer a more integrative characterization by enabling the simultaneous analysis of multiple genes. Nevertheless, these analyses still rely on extensive individual gene signatures, which may not accurately reflect actual alterations in the autophagic phenotype. Therefore, in this study, we established a functional gene set signature for autophagy monitoring that can indicate possible autophagy states. We demonstrated that its application in patient samples can be related to prognosis, mainly in glioblastoma. GBM is a highly aggressive tumor without an effective treatment strategy, so our characterization of autophagy at various levels for this cancer type adds new insights that contribute to GBM comprehension. Our results show that the induction state is more related to poor prognosis and comparisons between glioma grades suggest a negative correlation between lysosomal activity and glioma prognosis.

The separation between gene sets focused on the gene roles at each autophagic flux step and lysosome main elements, demanding an integrative analysis of multiple data sources. The resulting gene signature reflects the autophagy signaling pathway as intended, though it could be better optimized with the addition of more regulatory genes, such as other components of mTORC1 besides MTOR. In addition, concerning the usage of gene sets instead of individual genes, this approach seems to be highly effective considering the high regulatory aspects of the autophagy pathway. Mostly in cancer, alterations in specific ATGs, such as BECN1, ATG7 and ATG5, expression have already been shown though it seems to be cancer type dependent and differentially prognostic related [27–30]. However, individual gene alterations could not fully represent autophagy conditions, since consequent gene expression changes of functional homologs or related pathway components could compensate this individual alterations as already seen in other cases [31,32]. To minimize these effects and statistically enhance the analysis of differential expressions in gene sets between samples, we proposed the usage of GSVA methodology. This allows comparisons of gene set heterogeneity among samples, an essential element in cancer studies [23], and therefore an advantage of this approach in this study.

Regarding the autophagy states division, we started from the assumption that autophagy can be pharmacologically modulated and assume induction or inhibition profiles. However, even for data sets with the same pharmacological intervention, the differences among treatment conditions (time of exposure, drug concentration, time for RNA extraction, etc.) and sequencing methods caused a highly heterogeneous expression data between the analyzed series. It is important to consider that this is a usual challenge faced when dealing with high through output data [33]. These limitations led us to implement a supervised machine learning approach proposing two indexes—AutoIndex (AI) and LysoIndex (LI)—which, through a linear threshold (LI = 2AI – 0.2), allowed the separation of autophagy induction and inhibition states. These results reinforce the importance of lysosomal activity in characterizing autophagic flux functionality.

By applying this autophagic phenotypic 2D space to four cancer types — GBM, BRCA, LUAD, and COAD — we generated the overall hypothesis that tumors exhibit a higher incidence of an inhibition state compared to normal samples, although AI and LI means varied across cancer types. As described in the literature, we expected that high autophagic flux would be related to more aggressive tumors and poorer prognosis [10,11]. However, using our autophagy states division with the selected data, the gene set expression profiles indicated only a significative difference among states in GBM and less aggressive COAD samples, being the induction state more related to reduced survival. In addition, from all cancer types included in the analysis, GBM is the one with more significant influence of states division in clinical implications. Our findings suggest that the division among the induction–inhibition axis was much more important than the normal–tumor axis, indicating that low AI with high LI values tend to decrease survival and high AI with low LI, to increase. This relationship between GBM, lysosomal activity and prognosis is highlighted since LI values were only positive in GBM and only GBM showed an augmented and significant log2FC for a lysosomal gene set. Considering that GBM is the type of cancer with one of the highest malignancies [34], the presented results may indicate an association with high lysosomal activity and reduction of survival observed in GBM.

In order to verify if the alterations in lysosomal activity were not a particularity of the malignant neuronal tissue, we compared the previous GBM (G4) results with low glioma grades samples. LGG have a better prognosis when compared to GBM [25]. Differentially, G2 and G3 gliomas have lower LI values compared to G4 whether the alteration in AI was smaller. Moreover, lysosomal activity related gene sets showed a higher expression in G4, whereas autophagy regulation gene sets showed a lower expression when comparing different glioma grades. Besides that, when analyzing survival, G2 and G3 did not show significant differences between states, which can be related to the fact that these samples were much more shifted to high AI and low LI values. These results reinforce data already seen in the literature that indicates elevated lysosomal activity, indicated by high presence of lysosomal proteins like LAMP, in GBM [35,36].

Finally, we explored if the standard treatment for glioblastoma, TMZ and radiation therapy, could alter the states distribution. As previously seen in the literature, it is expected to observe a autophagy induction after TMZ treatment in glioblastomas [26]. With the expression data from patient-derived glioblastoma cell lines, we observed a tendency of treated samples, especially TMZ ones, to occupy the induction state. However, results were not statistically significant, possibly due to short samples sizes. In the glioma commercial cell lines data set, interestingly, treatment with TMZ significantly influenced one of the glioblastoma cell line (U87) occupancy of induction state, whereas the opposite was observed in the other (U251). These results suggest that autophagy alterations upon chemotherapy treatment could be cell type dependent. While the gene set showed relevance for GBM, it was less effective in representing other tumor types or lower grades. This suggests the current signature may be specific to more aggressive tumors, and a universal signature might require additional genes or data layers to capture broader tumor heterogeneity.

In summary, we proposed a functional autophagy gene set signature that considers both autophagic flux and lysosomal activity levels. With that, we established a gene set characterization method that can identify differential autophagy states in expression samples, categorizing them between induction and inhibition state based on AI and LI values. Applying this method to patient tumor samples, we showed a prevalence of inhibition state among GBM, BRCA, LUAD and COAD. Particularly, GBM was the cancer type with the highest correlation of *states separation axis* and patient survival. When compared to other glioma grades, the results indicate a possible relationship between GBM malignancy and higher LI values, with poorer prognosis being related to low AI values and high LI values. When exposed to TMZ treatment, patient-derived and commercial glioblastoma cell lines showed an increase of samples occupying the induction state, though further analysis with larger sample sizes are needed. These findings contribute to the comprehension and monitoring of autophagy and, specifically in GBM, point to a harmful role of autophagy, reinforcing its relevance as a therapeutic target.

## Materials and Methods

### Gene signature curation

The gene signature established for this analysis was defined by refined curation of open access repositories such as KEGG (https://www.kegg.jp/) and Reactome (https://reactome.org/) as well as related literature. The definition of genes related to autophagy was done by searching for mammal autophagy pathways at these databases and confirmed by the Human Autophagy Database (HADb, https://www.autophagy.lu/). The 81 chosen genes (Figure 1, Table S1) were divided into 9 gene sets (Initiation, Nucleation, Elongation, Maturation, Positive Regulation, Negative Regulation, Lysosomal Transporters, Lysosomal Enzymes and Lysosomal Membrane). There were also defined 2 gene set groups: Autophagy (Initiation, Nucleation, Elongation and Maturation) and Lysosomal Activity (Lysosomal Transporters, Lysosomal Enzymes and Lysosomal Membrane).

### Autophagy modulation data sets and states defining

#### Data origin

Selection of data sets with autophagy modulation in cancer or normal cell lines was done among GEO (https://www.ncbi.nlm.nih.gov/geo/) public data. There were 19 data sets selected (Table 1, Table S2): 12 with autophagy induced conditions (GSE276412, GSE130006, GSE242110, GSE99875, GSE202227, GSE196231, GSE218784, GSE252314, GSE246170, GSE271357, GSE180321) and 8 with autophagy inhibited conditions (GSE154936, GSE128844, GSE141507, GSE60570, GSE90444, GSE217572, GSE171167, GSE255564). Every data set contained bulk RNA-seq data obtained by high through output sequencing and both control and treated samples. All expression data was normalized to TPM, if not already as TPM available, and filtered for the gene signature previously defined.

#### Differential gene (set) expression analysis

The gene set analysis was done using the R-package GSVA (Gene Set Variance Analysis, https://bioconductor.org/packages/release/bioc/html/GSVA.html). For every gene set in each sample, there was established a GSVA score with which was done differential gene set expression analysis with R-package limma (https://bioconductor.org/packages/release/bioc/html/limma.html), as recommended by the package guidelines. The default parameters were set, and the lof2FC values were calculated comparing treated and control samples. For the differential gene expression analysis of specific autophagy markers [15], the DEGs were defined using limma package with |log2FC|>1.0 and p-value<0.05.

#### States clustering

The log2FC change values of each data set were used for two types of clusterization: heatmap and a supervised machine learning approach. The heatmap was generated with R-package pheatmap (https://cran.r-project.org/web/packages/pheatmap/index.html) and both columns and rows were clustered with default parameters (distance euclidean and method complete). The supervised machine learning approach was done with a Visual Basic pipeline. The separation threshold was defined to better group samples of the same condition (induction or inhibition). For each gene set was established a coefficient value which was used to define two index values: AutoIndex (Figure 3D) and LysoIndex (Figure 3E). The separation line is *y = 2x - 0.2*, where *y* stays for LysoIndex and *x* for AutoIndex. The automated table for index calculation and phenotypic space plotting can be found in Table S3.

### Clinical pan-cancer data

#### Tumor samples

Patient derived tumor data was obtained from TCGA (https://www.cancer.gov/ccg/research/genome-sequencing/tcga) for GBM (Glioblastoma Multiforme), BRCA (Breast Invasive Carcinoma), LUAD (Lung Adenocarcinoma) and COAD (Colon Adenocarcinoma) projects (Table S4). The data were directly in the R terminal downloaded with the package TCGAbiolinks (https://bioconductor.org/packages/release/bioc/html/ TCGAbiolinks.html). Bulk RNA-seq data were obtained with filters “experimental.strategy = RNA-Seq”, “data.category = Transcriptome Profiling”, “data.type = Gene Expression Quantification” and “workflow.type = STAR – Counts”. Clinical data was directly obtained with function “GDCquery_clinic()”. The tumor samples were filtered for “Primary Solid Tumor” and randomly sampled for 367 samples per tumor type (maximal Tumor samples of GBM available). Expression data were normalized into TPM and filtered for the gene signature before expression analysis.

#### Normal samples

Normal tissue samples were obtained directly from the website of The Adult Genotype Tissue Expression Project (GTEx). There were selected 367 normal samples for each tumor type between 1 or 2 tissues related (Table S5). Normal brain samples were selected among cortex and frontal cortex tissues, normal breast samples among mammary tissue, normal lung samples among lung tissue and normal colon samples among Colon Sigmoid and Colon Transverse tissues. Expression data were normalized into TPM and filtered for the gene signature before expression analysis.

#### States analysis

For each gene set of each sample a GSVA score was calculated. Log2FC values were obtained for the whole cohort (Tumor x Control) and for each sample (mean expression of control samples was used as denominator group). Cohort log2FC values were displayed as a volcano plot generated with R-package EnhancedVolcano (https://bioconductor.org/packages/release/bioc/html/EnhancedVolcano.html). Individual log2FC values were used to calculate Auto and LysoIndex. Dispersion values for each individual index were also calculated.

#### Survival analysis

Survival data was obtained from clinical data of the TCGA projects and overall survival was calculated in days. Kaplan-Meier analysis was performed for each tumor type, using the median to separate patients with an autophagy induction and inhibition state. In addition to the main analysis, three complementary analyses were performed independently: (i) filtering for patients with overall survival greater than 300 days; (ii) dividing patients into two groups based on the median transcriptomic distance to normal samples (closer 50% vs. farther 50%); and (iii) stratifying patients into tertiles according to the states division. The Kaplan-Meier curves were generated with Cox Regression at GraphPad Prism 8. P-values <0.05 were statistically significant.

### Other glioblastoma conditions and treatment comparisons

Low glioma grade data were obtained from the TCGA Project LGG (Table S4). The bulk RNA-seq data and clinical information were downloaded as previously described for other projects. There were selected 200 samples from each glioma grade available (G2 and G3) and 200 normal samples were selected from the previous GBM cohort of normal brain samples. For treatment comparisons, there were selected 2 GEO datasets (Table S6): GSE249544 (patient-derived glioblastoma cell lines exposed to radiation or temozolomide) and GSE229600 (glioblastoma and astrocytoma cell lines exposed to temozolomide). Each data set contained treated and control conditions. The expression data was used to analyze states distributions upon treatment as previously described.

### Statistical analysis

Statistical analyses were carried out using GraphPad Prism 8. Normality was tested with Shapiro–Wilk test and following statistical analysis was performed for two groups comparisons either with Mann–Whitney U test (no Gaussian distribution) or unpaired two-tailed t-test (Gaussian distribution), or multiple comparisons either with or Kruskal-Wallis test followed by Dunn’s test (no Gaussian distribution) or ANOVA followed by Tukey’s post-hoc test (Gaussian distribution). P-values <0.05 were statistically significant.

## Abbreviations

PAS: Phagophore assembly site
PG: phagophore
ATG: autophagy related gene
HADb: Human Autophagy Database
HAMdb: Human Autophagy Modulator Database
GSVA: Gene Set Variation Analysis
TCGA: The Cancer Genome Atlas
GBM: Glioblastoma Multiforme
BRCA: Breast Invasive Carcinoma
LUAD: Lung Adenocarcinoma
COAD: Colon Adenocarcinoma
LGG: Low Grade Glioma
ARG: autophagy related gene
KEGG: Kyoto Encyclopedia of Genes and Genomes
L. activity: Lysosomal activity
L. membrane: Lysosomal membrane
L. transporters: Lysosomal transporters
L. enzymes: Lysosomal enzymes
GEO: Gene Expression Omnibus
TPM: transcripts per million
log2FC: log2 fold change
ML: machine learning
AI: AutoIndex
LI: LysoIndex
TMZ: temozolomide
GSEA: Gene Set Enrichment Analysis
GTEx: The Adult Genotype Tissue Expression Project

## Data availability

The datasets for autophagy states definition and glioma treatment comparisons can be found in GEO repository under the accession numbers included at Table S2 and Table S6, respectively. Clinical pan-cancer data are available in the GDC Data Portal. The code to reproduce the analyses and some of the figures described in this study can be found at https://github.com/camillabm/TCC.

## Author contributions

CRediT: **Camilla Brückmann de Mattos:** Conceptualization, Formal Analysis, Investigation, Data curation, Visualization, Writing – original draft. **Daphne Torgo:** Conceptualization, Supervision, Writing – review & editing. **Guido Lenz:** Conceptualization, Supervision, Writing – review & editing.

## References

[1] Glick D, Barth S, Macleod KF. Autophagy: cellular and molecular mechanisms. J Pathol. 2010;221(1):3–12.

[2] Yang Z, Klionsky DJ. Mammalian autophagy: core molecular machinery and signaling regulation. Curr Opin Cell Biol. 2010;22(2):124–131.

[3] Parzych KR, Klionsky DJ. An Overview of Autophagy: Morphology, Mechanism, and Regulation. Antioxid Redox Signal. 2014;20(3):460–473.

[4] Torres-López L, Dobrovinskaya O. Dissecting the Role of Autophagy-Related Proteins in Cancer Metabolism and Plasticity. Cells. 2023;12(20):2486.

[5] Russell RC, Yuan H-X, Guan K-L. Autophagy regulation by nutrient signaling. Cell Res. 2014;24(1):42–57.

[6] Levine B, Kroemer G. Autophagy in the Pathogenesis of Disease. Cell. 2008;132(1):27–42.

[7] Mizushima N, Levine B, Cuervo AM, et al. Autophagy fights disease through cellular self-digestion. Nature. 2008;451(7182):1069–1075.

[8] Mulcahy Levy JM, Thorburn A. Autophagy in cancer: moving from understanding mechanism to improving therapy responses in patients. Cell Death Differ. 2020;27(3):843–857.

[9] Li X, He S, Ma B. Autophagy and autophagy-related proteins in cancer. Mol Cancer. 2020;19(1):12.

[10] Ma X-H, Piao S, Wang D, et al. Measurements of tumor cell autophagy predict invasiveness, resistance to chemotherapy, and survival in melanoma. Clin Cancer Res Off J Am Assoc Cancer Res. 2011;17(10):3478–3489.

[11] Koh M, Lim, Hyesol, Jin, Hao, et al. ANXA2 (annexin A2) is crucial to ATG7-mediated autophagy, leading to tumor aggressiveness in triple-negative breast cancer cells. Autophagy. 2024;20(3):659–674.

[12] Carew JS, Nawrocki ST, Kahue CN, et al. Targeting autophagy augments the anticancer activity of the histone deacetylase inhibitor SAHA to overcome Bcr-Abl– mediated drug resistance. Blood. 2007;110(1):313–322.

[13] Amaravadi RK, Yu D, Lum JJ, et al. Autophagy inhibition enhances therapy-induced apoptosis in a Myc-induced model of lymphoma. J Clin Invest. 2007;117(2):326–336.

[14] Loos B, du Toit A, Hofmeyr J-HS. Defining and measuring autophagosome flux— concept and reality. Autophagy. 2014;10(11):2087–2096.

[15] Klionsky DJ, Abdel-Aziz, Amal Kamal, Abdelfatah, Sara, et al. Guidelines for the use and interpretation of assays for monitoring autophagy (4th edition)1. Autophagy. 2021;17(1):1–382.

[16] Kim J, Eberwine J. RNA: State Memory and Mediator of Cellular Phenotype. Trends Cell Biol. 2010;20(6):311–318.

[17] Deng J, Zhang Q, Lv L, et al. Identification of an autophagy-related gene signature for predicting prognosis and immune activity in pancreatic adenocarcinoma. Sci Rep. 2022;12(1):7006.

[18] Lin X, Wu Q, Qin S, et al. Identification of an Autophagy-Related Gene Signature for the Prediction of Prognosis in Early-Stage Colorectal Cancer. Front Genet [Internet]. 2021 [cited 2025 May 22];12.

[19] Bordi M, De Cegli R, Testa B, et al. A gene toolbox for monitoring autophagy transcription. Cell Death Dis. 2021;12(11):1–7.

[20] Sager R. Expression genetics in cancer: Shifting the focus from DNA to RNA. Proc Natl Acad Sci U S A. 1997;94(3):952–955.

[21] Hale AN, Ledbetter DJ, Gawriluk TR, et al. Autophagy. Autophagy. 2013;9(7):951–972.

[22] Galluzzi L, Bravo-San Pedro JM, Levine B, et al. Pharmacological modulation of autophagy: therapeutic potential and persisting obstacles. Nat Rev Drug Discov. 2017;16(7):487–511.

[23] Lenz G, Onzi GR, Lenz LS, et al. The Origins of Phenotypic Heterogeneity in Cancer. Cancer Res. 2022;82(1):3–11.

[24] Becker AP, Sells BE, Haque SJ, et al. Tumor Heterogeneity in Glioblastomas: From Light Microscopy to Molecular Pathology. Cancers. 2021;13(4):761.

[25] Li D, Chen Y, Wong T-F, et al. Management and survival trends for diffuse gliomas diagnosed at a single neurooncology center in China during 2000 to 2020. Sci Rep. 2025;15(1):12574.

[26] Würstle S, Schneider F, Ringel F, et al. Temozolomide induces autophagy in primary and established glioblastoma cells in an EGFR independent manner. Oncol Lett. 2017;14(1):322–328.

[27] Liang XH, Jackson S, Seaman M, et al. Induction of autophagy and inhibition of tumorigenesis by beclin 1. Nature. 1999;402(6762):672–676.

[28] Cao Q-H, Liu F, Yang Z-L, et al. Prognostic value of autophagy related proteins ULK1, Beclin 1, ATG3, ATG5, ATG7, ATG9, ATG10, ATG12, LC3B and p62/SQSTM1 in gastric cancer. Am J Transl Res. 2016;8(9):3831–3847.

[29] Zhu J, Li Y, Tian Z, et al. ATG7 Overexpression Is Crucial for Tumorigenic Growth of Bladder Cancer In Vitro and In Vivo by Targeting the ETS2/miRNA196b/FOXO1/p27 Axis. Mol Ther Nucleic Acids. 2017;7:299–313.

[30] Ge J, Chen Z, Huang J, et al. Upregulation of autophagy-related gene-5 (ATG-5) is associated with chemoresistance in human gastric cancer. PloS One. 2014;9(10):e110293.

[31] Chang Y, Lu X, Qiu J. Compensatory expression regulation of highly homologous proteins HNRNPA1 and HNRNPA2. Turk J Biol Turk Biyol Derg. 2021;45(2):187–195.

[32] Barth AS, Kumordzie A, Frangakis C, et al. Reciprocal Transcriptional Regulation of Metabolic and Signaling Pathways Correlates With Disease Severity in Heart Failure. Circ Cardiovasc Genet. 2011;4(5):475–483.

[33] Yu Y, Mai Y, Zheng Y, et al. Assessing and mitigating batch effects in large-scale omics studies. Genome Biol. 2024;25(1):254.

[34] Tran B, Rosenthal MA. Survival comparison between glioblastoma multiforme and other incurable cancers. J Clin Neurosci. 2010;17(4):417–421.

[35] Jensen SS, Aaberg-Jessen C, Christensen KG, et al. Expression of the lysosomal-associated membrane protein-1 (LAMP-1) in astrocytomas. Int J Clin Exp Pathol. 2013;6(7):1294–1305.

[36] Sarafian VS, Koev I, Mehterov N, et al. LAMP-1 gene is overexpressed in high grade glioma. APMIS Acta Pathol Microbiol Immunol Scand. 2018;126(8):657–662.

