## Supplementary figures for "Functional autophagy gene set signature and state classification reveal a link between autophagy induction, lysosomal activity, and poor prognosis in glioblastoma"

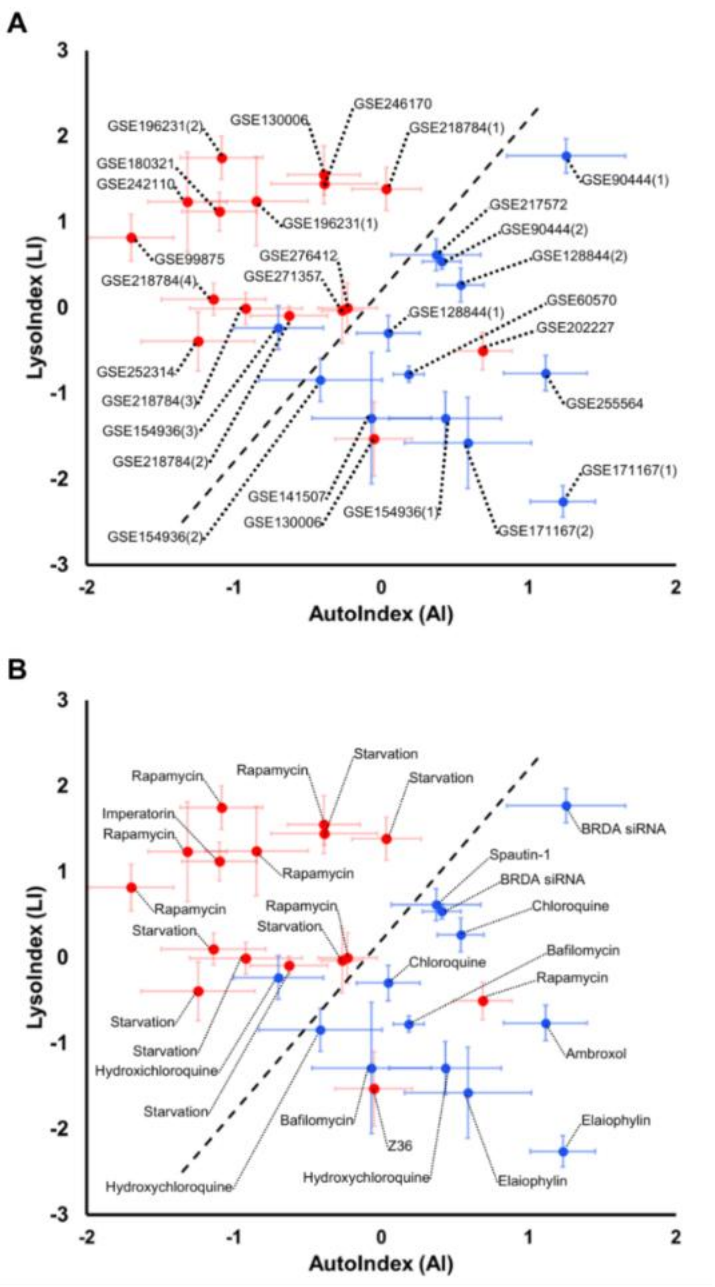


**Figure S1.** GEO autophagy states phenotypic states plot. (**A**) Phenotypic states plot labeled with GEO Series. (**B**) Phenotypic states plot labeled with autophagy modulator used.

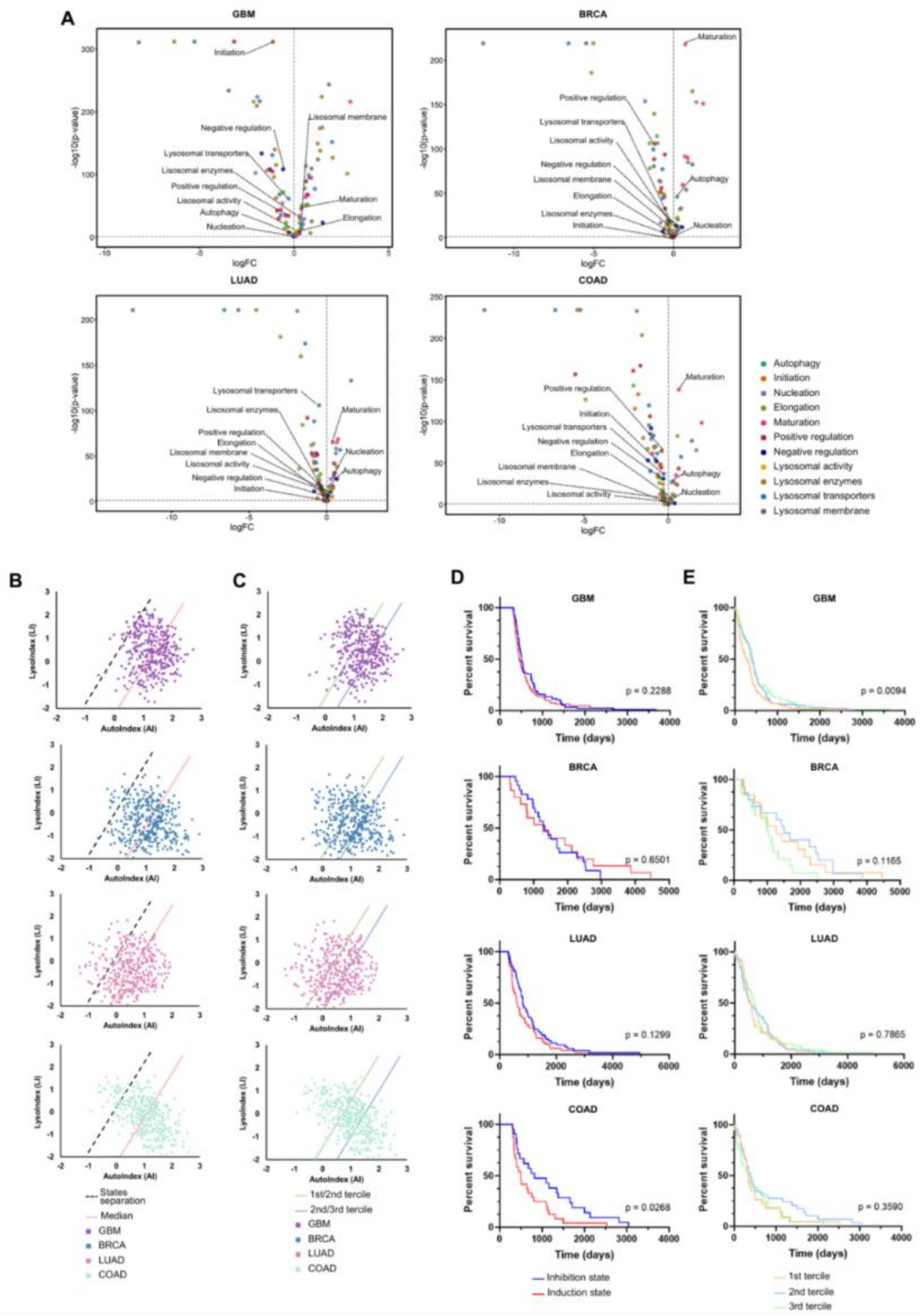

**Figure S2.** Complementary data for autophagy states characterization among cancer types. (**A**) Volcano plots with color differentiation of gene sets and related genes. The ones labelled are points that indicate log2FC and p-values for the gene sets. (**B**) Tumor samples divided into the median. (**C**) Tumor samples divided into terciles.

Kaplan-Meier survival analysis for each cancer type comparing autophagy states for samples with at least 300 days of overall survival (**D**) and samples divided into terciles (**E**). P-values are indicated.

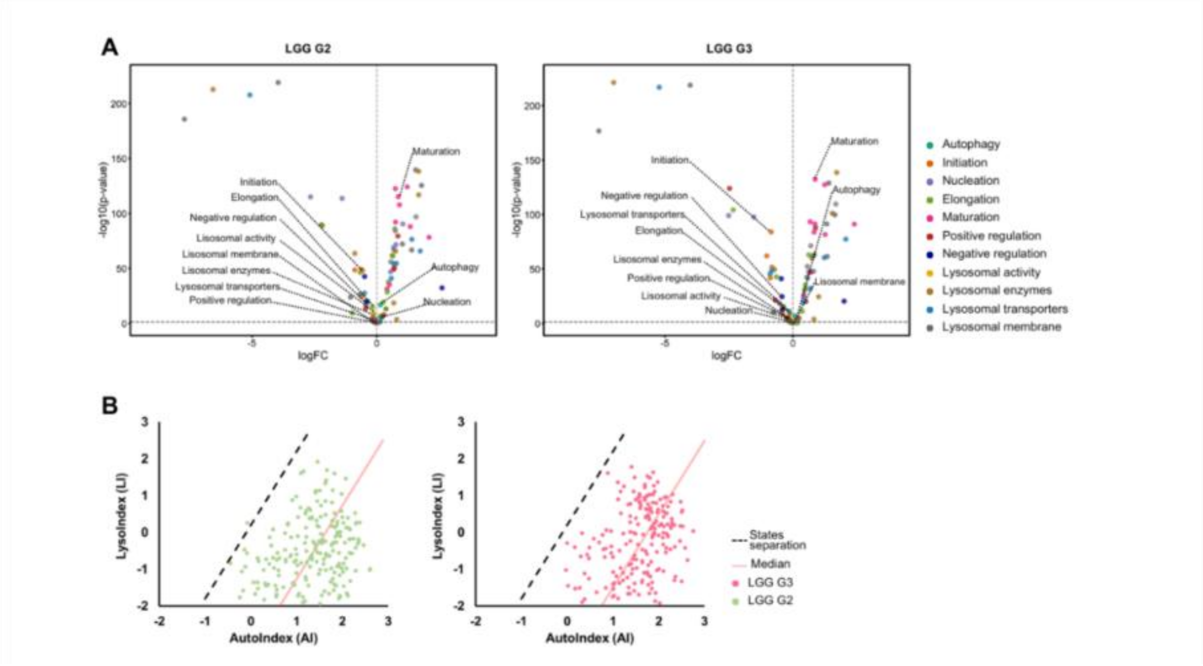


**Figure S3.** Complementary data for autophagy states characterization among glioma grades. (**A**) Volcano plots with color differentiation of gene sets and related genes. The ones labelled are points that indicate log2FC and p-values for the gene sets. (**B**) Tumor samples divided into the median.
